# Divergent molecular and circuit mechanisms underlie light entrainment of retinal and suprachiasmatic nucleus circadian clocks

**DOI:** 10.64898/2026.04.27.721075

**Authors:** A Jandot, H Calligaro, A Dianak-Shoori, K Tartour, C Sandu, N Haddjeri, CP Ribelayga, B Ananthasubramaniam, K Padmanabhan, MP Felder-Schmittbuhl, O Dkhissi-Benyahya

## Abstract

The circadian system aligns behavior and physiology with the 24-hour environmental cycle through a distributed network of clocks including the master pacemaker in the suprachiasmatic nucleus (SCN) and an autonomous retinal clock critical for local retinal physiology and function. Although both clocks are entrained by light, they differ in their photoreceptor inputs and light sensitivity. The specific contributions and mechanisms by which distinct photoreceptor pathways drive their photoentrainment, however, remain incompletely understood.

In this study, we conducted a comprehensive transcriptomic and integrative comparative analysis of retinal and SCN circadian light responses using mouse models lacking specific photoreceptors or key components of signaling pathways. Under photopic conditions, we found that each tissue displays distinct light-responsive transcriptional signatures across genotypes, yet both shared a conserved cluster of rod-driven immediate early-genes. Strikingly, the light-evoked transcriptional response was not sufficient to shift the phase of the SCN clock, in contrast to its robust phase-shifting effect on the retinal clock. Furthermore, by genetically disrupting rod/cone electrical coupling and pharmacologically isolating rod pathways, we identified the OFF-cone bipolar cell circuit as both necessary and sufficient to mediate light-induced phase resetting of the retinal clock. Together, these findings delineate the specialized retinal circuitry that underlies circadian entrainment and reveal a fundamental divergence between retinal and SCN mechanisms of photic timekeeping.

## Introduction

Living organisms rely on endogenous circadian clocks to synchronize their behavior and physiology with the environmental light/dark (LD) cycle. In mammals, the primary external time cue is environmental light, which can vary in intensity by up to a billion-fold between night and day [1]. Light is received by retinal photoreceptors, including rods, cones, and intrinsically photosensitive retinal ganglion cells (ipRGCs), all of which contribute to non-image-forming responses such as the pupillary light reflex and the circadian photoentrainment of the master clock, located in the suprachiasmatic nuclei (SCN) of the hypothalamus [2–10]. Beyond entraining the SCN, the mammalian retina harbors an autonomous circadian clock [11] that functions independently of SCN control [12]. The retinal clock orchestrates critical retinal functions [13] including photoreceptor disc shedding [14–17], melatonin and dopamine release [18,19], photoreceptor electrical coupling [20–22], the amplitude of light responses [23,24], circadian clock gene expression [25,26] and visual processing [12].

Although light-induced phase shifts of the SCN clock, driven by the rapid and transient induction of immediate-early genes such as *Fos*, are well characterized [27–32], direct comparative analyses of light-evoked responses between retinal and SCN clocks remain limited. Our previous work showed that the retina exhibits a markedly lower threshold for light-induced FOS expression, approximately 1-2 log units below that of the SCN [33]. Intringuinly, phase shifting of the retinal clock requires substantially higher irradiance (∼10^13^ photons/cm^2^/s [34], than that required to elicit behavioral phase shifts (10^10^-10^11^ photons/cm^2^/s, [4,5,34–36]). We recently demonstrated that, under photopic conditions, rods, rather than middle-wavelength (MW) cones or melanospin, are essential for light-induced phase shifting of the mouse retinal clock but not SCN enterainment [34,37]. The SCN receives photic input exclusively via ipRGCs, yet these cells integrate rod-driven signals to mediate circadian entrainment [2,5,6].

The broad dynamic range of rod involvement across varying light intensities can be attributed to the engagement of ON and OFF pathways through three distinct rod-mediated circuits, each characterized by different sensitivities [38]. The primary pathway is the most sensitive, operating near scotopic threshold. It transmits light signals from rods to ON-type rod bipolar cells (ON-RBCs) expressing metabotropic glutamate receptor 6 (mGluR6) [38,39], which then synapses onto AII amacrine cells. These amacrine cells connect to ON-type cone bipolar cells (ON-CBCs) *via* gap junctions and to OFF-CBCs through inhibitory glycinergic synapses, ultimately relaying signals to ON and OFF retinal ganglion cells (RGCs), respectively. A secondary pathway, recruited at higher light levels, conveys rod signals directly to adjacent cones through connexin 36 (*Cx36*) gap junctions. When rods hyperpolarize, glutamate release ceases, leading to depolarization of OFF-CBCs via ionotropic kainate/AMPA-type glutamate receptors. Conversely, ON-CBCs hyperpolarize through activation of mGluR6 receptors, resulting in reduced cationic conductance [40,41]. These CBCs then convey the signal to RGCs [42–44]. The third and least characterized pathway involves direct synaptic rod input to OFF-CBCs [44–47]. Despite the delineation of these circuits, the specific rod pathway responsible for mediating light-induced phase shifts of the retinal clock remains unknown.

To isolate rod-specific transcriptomic signatures elicited by light in the SCN and retina, and to dissect the contribution of individual rod pathways to retinal clock phase shifting, we first investigated light-driven gene activation in isolated retinal layers and in the SCN of the same wild-type and “rod-only” mice. Transcriptomic profiling revealed tissue-specific light-responsive gene expression in both genotypes, along with a shared cluster of immediate early-genes. To further characterize these responses, we assessed FOS protein induction, a canonical marker of photic activation across tissues. We found that rod loss differentially impaired FOS expression and phase-shifting responses in the retina and SCN. Finally, we identified the OFF-CBC pathway as the principal route through which rods mediate light-induced phase shifts of the retinal clock. Together, these findings provide compelling evidence that, under photopic conditions, the retinal and SCN clocks rely on distinct photoreceptors and downstream signaling pathways to process environmental light cues.

## Materials and Methods

### Animals

Mice were housed in temperature-controlled room (23 ± 1°C), under 12 h light/12 h dark cycle (12L/12D, light intensity around 200 lux) with food and water *ad libitum*. All animal procedures were in strict accordance with current national and international regulations on animal care, housing, breeding and experimentation and approved by the regional ethics committee CELYNE (C2EA42-13-02-0402-005). All efforts were made to minimize suffering. *Per2^Luc^* mice [48] and several photoreceptor deficient mice were used: *TRβ ^-/-^*lacking MW cones [3,49], *Opn4^-/-^* knockout for melanopsin [5] and *Nrl^-/-^* characterized by the complete loss of rods and an increased number of SW cones [50,51], as well as mice knocked out for connexin 36 either globally (*Cx36^-/-^*) or specifically in rods (*Rhoi75^Cre^::Cx36^flox/flox^*) or cones (*HRGP^Cre^::Cx36^flox/flox^*) [52,53]. All mice with the exception of *HRGP^Cre^::Cx36^flox/flox^* were bred with the *Per2^Luc^* mice to obtain *Opn ^-/-^::TRβ ^-/-^::Per2^Luc^*, *Nrl^-/-^::Per2^Luc^*, *Cx36^-/-^::Per2^Luc^* and *Rhoi75^Cre^::Cx36^flox/flox^::Per2^Luc^*. All lines were maintained on a C57BL/6J background. We used female and male mice in all experiments, except for behavioral phase shifts where only males were used to avoid confounding parameters [54]. All mice were used between 2-4 month-old, except for the *Nrl^-/-^::Per2^Luc^* model in culture which were used at 1 month, before the onset of apoptotic degeneration [55]. *Nrl^- /-^* mice were also used at 5 months in behavior and immunolabelling experiments to avoid noise and immunoreactivity independent of light caused by the outer retina degeneration.

### Laser capture microdissection of the retinal layers and RNA extraction

Wild-type (n=9) and *Opn ^-/-^::TRβ ^-/-^* mice (n=6) were exposed to a single pulse of monochromatic light (530 nm) of constant duration and irradiance (30 min, 10^14^ photons/cm^2^/s) at CT16 (first day in DD). Animals were sacrificed and the SCN and both eyes from the same animal were dissected at CT16 (DC), 30 min or 2h (only for WT mice) after the beginning of the light pulse, rapidly frozen on dry ice and stored at -80°C. All dissections during the dark phase were carried out under dim red light. Frozen eyes were cut at 12 µm thickness and mounted on Glass PEN membrane slides (Leica). The frozen sections were thawed for 30 s and were immersed immediately in ethanol solutions (95%, 75%, 50%) for 30s each. Few drops of 1% Cresyl violet acetate in 50% ethanol were applied to the sections for 15 s. The slides were then rinsed in 2 successive baths of 50% ethanol, dehydrated in graded ethanol baths (75%, 95%, 100% twice) for 30 s each and air-dried in a vacuum desiccator for a minimum of 5 min. All the ethanol baths contain 0.2% ProtectRNA RNase inhibitor 500x (Sigma) except for the last one. In order to isolate the retinal layers in a contact- and contamination-free manner, the Leica LMD6 laser microdissection was used. Laser capture was performed by lifting separately these 3 layers by gravity into the cap of an Eppendorf tube filled with 20 μl buffer RLT (Qiagen Rneasy Micro Kit) including 2M DTT. RNAs were extracted using Trizol reagent (Invitrogen) and their qualities assessed on an Agilent 2100 Bioanalyzer (Agilent Technologies, USA).

### RNA sequencing and Gene ontology analysis

RNASeq Total RNA was prepared using the “Stranded Total RNA Prep with RiboZero” kit (Illumina), following the manufacturer’s instructions to deplete ribosomal RNA and enrich for coding and non-coding RNA species. Sequencing was performed on 24 samples using the Illumina NextSeq 500 platform with the High Output kit, generating single-end 76 bp reads and dual indexing (8 bp i7 + 8 bp i5). A total of over 526 million reads were obtained, ranging from 17 to 25 million reads per sample with % Q30 >89%. Raw RNA-seq reads in FASTQ format were quantified using Salmon (v.1.10.3) [56] in default mode. Indexing was performed using the salmon index command on the mouse transcriptome (GENCODE release M36). Transcript-level quantifications from Salmon were imported into R (v.4.4.2) using the tximport (v.1.34.0) package [57] and summarized to gene-level counts using a transcript-to-gene mapping table derived from the comprehensive gene annotation GTF file (GRCm39). The gene-level counts were used for downstream differential expression analysis with DESeq2 (v.1.46.0) [58] to estimate absolute fold-changes and P-values. P-values were adjusted for false discovery rate (FDR) control using the Benjamini-Hochberg correction. Differentially expressed genes were defined as those with FDR < 0.05 and |log2(fold-change)| > 1.0. Comparison between 30 minutes light exposure and their respective dark controls in WT and *Opn4^-/-^::Trβ ^-/-^* samples in different retinal layers (ONL, INL and GCL) and the SCN was performed with appropriate contrasts and visualized with volcano plots. These genes were plotted via VennDiagram (v.1.7.3) [59] to visualize the number of genes in common among WT and the “rod-only” mutant in each layer. Genes were initially hierarchically clustered using base functions, grounded on the similarity of their light response (absolute log2 fold-change of differential gene expression) across retinal layers and genotypes using Euclidean distance on scaled expression values as the metric. From the heatmap several clusters were selected based on biological relevance and similarity of light response.

### Functional enrichment

Gene IDs of differentially expressed genes were mapped by the means of org.Mm.eg.db [60] package. The mapped IDs were the subject of functional annotation enrichment analysis using clusterProfiler (v.4.14.6) [61] package. Significant genes were filtered for adjusted p-value < 0.05 plus |log2(fold-change)| > 1.0 as before. Number of significant upregulated and downregulated genes were determined with thresholds set such as previously to adjusted p-value < 0.05 plus |log2(fold-change)| > 1.0 as before for over-representation analysis (which, is included in our results as the plot of gene ontology (GO) enriched terms) in addition to the mapped gene set enriched terms (GSEA) (which, their results is presented as the map of enriched terms from Kyoto encyclopedia of genes and genomes (KEGG)) based on all the genes included in the data set ranked by a decreasing absolute of fold-change value.

### Retinal explant culture and bioluminescence recording

Mice were killed by cervical dislocation 1 h before light offset (Zeitgeber Time 11 or ZT11). Eyes were enucleated and placed in Hank’s balanced salt solution (HBSS; Invitrogen). Retinas were gently isolated from the rest of the eye cup and flattened ganglion cell layer up on a semi-permeable (Millicell) membrane in a 35 mm culture dish (Nunclon) containing 1.2 mL Neurobasal-A (Life Technologies) with 2% B27 (Gibco), 2 mM L-Glutamin (Life Technologies) and 25 U/mL antibiotics (Penicilin/Streptomycin, Sigma), incubated at 37°C in 5% CO_2_ for 24 h. From this step on, all manipulations of explants were performed under dim red light. After 24h, at the projected ZT12, retinas were transferred to 1.2 ml of 199 medium (Sigma), supplemented by 4 mM sodium bicarbonate (Sigma), 20 mM D-glucose (Sigma), 2% B27, 0.7 mM L-Glutamin, 25 U/mL antibiotics (Penicilin / Streptomycin, Sigma) and 0.1 mM Luciferin (Perkin). Culture dishes were sealed and then placed in a Lumicycle (Actimetrics, Wilmette, IL, USA) to record the global emitted bioluminescence. PER2::Luc bioluminescence was analysed using Lumicycle Analysis software (Actimetrics, Wilmette, IL, USA). Bioluminescence data were detrended using a 25 h running average and the phase and the period were determined by using best-fit sine wave function of the Lumicycle Analysis software. To analyse potential genotype-related differences, we evaluated whether the endogenous functioning of the retinal and SCN clocks was altered across the different genotypes. We found a shortened period of the retinal clock and of the rhythm of behavior in “rod-only” and *Rod^Cre^::Cx36^fl/fl^* mice by comparison to WT (p<0.001). Interestingly, the endogenous period of PER2::Luc oscillations was significantly lengthened in *Cx36^-/-^* retinal explants (25.84 ± 0.2h, p<0.001), suggesting a critical role for *Cx36*-mediated coupling in the inner retina to maintain synchrony among retinal oscillators. A similar trend was observed for the endogenous period calculated for the rhythm of locomotor behavior.

### Phase shifting response of the retina clock and data analysis

Light stimulations were done at CT16 as previously described [34,37]. We used constant irradiance and duration (10^14^ photons/cm²/s, 30 min) of a monochromatic light (520 nm). Since the retinal pigment epithelium was removed to prevent confounding effects from its rhythmic PER2::Luc expression, we applied only a single light stimulus to each retinal explant in order to avoid any photopigment depletion. Phase shifts are calculated as the difference between the predicted phase and the measured peak of PER2::Luc, corrected by the endogenous period measured on the 3 days before light stimulation. Endogenous and light-affected period were determined using 3 oscillations either before or after light stimulation respectively. Light wavelength and intensity were measured using a spectrophotometer (ILT 5000, International Light Technologies). To isolate the specific effect of light on the phase, independent of its effects on period, or potential genotype-related differences, we compared the impact of light on the endogenous period among genotypes (difference between the “period after” minus “period before” the light stimulation). We found a similar lengthening of the period in WT, “rodless” and “rod-only” retinal explants after the light pulse while light shorthen the period in *Cx36^-/-^* explants and did not change the period in *Rod^Cre^::Cx36^fl/fl^* (data not shown). For the behavioral phase shift, light caused a lengthening of the period in all genotypes, except for “rod-only” mice (data not shown). The effect of light on period have been taken into account for the phase shift calculation for both the retina and the SCN.

### Retinal explant culture and pharmacological treatments

The 199 medium was supplemented with L-AP4 (5µM) to block mGluR6 receptors of the ON pathway or with a combinaison of L-AP4 and UBP-310 (50µM) to additionally block AMPA/kainate receptors of the OFF pathways. The impact of the these pharmacological agents on the endogenous period and phase of the retinal clock were examined. We observed a lengthening of the endogenous period and a delay in the phase of PER2::LUC oscillations when retinal explants are incubated with L-AP4 (25.27 ± 0.17h; CT 23.8 ± 0.33; p<0.001, data not shown) by comparison to non-supplemented retinas (24.68 ± 0.07h; CT21.91 ± 0.46). To ensure accurate assessment of light-induced phase shifts, values were calculated relative to a baseline specific to each explant, thereby controlling for initial phase differences between conditions and samples. Additionally, all measurements were corrected for period length changes induced by light and experimental treatment, following the approach described by [34,37].

### Behavioural Phase-Shifting Assay

Singly housed male WT, *Nrl^-/-^*, *Opn ^-/-^::TRβ ^-/-^*, *Cx36^-/-,^ Rhoi75^Cre^::Cx36^flox/flox^* and *HRGP^Cre^::Cx36^flox/flox^* mice (WT : n = 10 ; *Nrl^-/-^*: n=8 ; *Opn ^-/-^::TRβ ^-/-^* n = 10 ; *Cx36^-/-^* n = 7 ; *Rhoi75^Cre^::Cx36^flox/flox^* n = 10 and *HRGP^Cre^::Cx36^flox/flox^* n= 5) were first entrained in a 12L/12D cycle for 15 to 20 days. Subsequently, animals were maintained in constant darkness (DD) to examine the free-running period calculated by periodogram analysis using ClockLab software (Actimetrics). Phase shifts were studied using a 30 min monochromatic light pulse (530 nm, half-bandwidth, 10 nm) at 10^14^ photons/cm²/s applied at CT16. The stimulator (light source and chamber) has been described previously [33]. After the light pulse, animals were returned to their home cages, and activity was monitored in DD for an additional 10 days before the next light pulse. The magnitude of a light-induced phase shift was determined from the difference between the regression lines of the activity onsets before and after the light stimulation, extrapolated to the day following the light pulse. The transient responses on the 2 days immediately after the pulse were discounted for the determination of activity onsets.

### FOS immunohistochemistry

Mice were kept in dim red light until the end of the fixation. 1h after the beginning of the light stimulation, mice were anaesthetized with 2.5µL/g ketamine and 5µL/g sodium pentobarbital, and perfused with a 4% paraformaldehyde in 0.1M phosphate buffer (PB) solution. Eyes and brain are collected, rinsed in PB and cryoprotected with 30% sucrose solution. Eyes are cut in 100 sagittal sections of 20µm from the point where the 3 layers of the retina can be separated and collected on 25 serialized slides of 4 sections each. Brains are cut in 30 serialized rostral sections of 35µm from the beginning of the SCN, localized on top of the optic chiasma. One every 3 sections was stained and quantified in the SCN; 1 every 5 slides were stained for the retina, and 1 out of 2 consecutive stained sections was quantified to avoid double counting of a cell and cover the largest area in a sample. Sections were incubated overnight with the FOS rat antibody (Ab 226 017, Synaptic Systems, 1/10 000), then incubated with biotinylated rabbit anti-rat antibody for 2 h (BA 4000 biotinylated, Vector, 1/200), streptavidin-Cy3 for 2 h (Jackson, 1/500), and DAPI for 5 min (Jackson, 1/10,000). Image for quantification were obtained every 6 µm using Zeiss Slide Scanner to build a z-projection of the section and reveal all labelled cells in the tissue depth. Quantification was done using Neurolucida software (MBF Biosciences) by defining thresholds in labelling intensity and cell size for each area of interest. Micrograph were obtained with Leica confocal microscopy.

### Statistical analysis

Statistical analyzes were performed using two-way ANOVA followed, when significant (p ≤ 0.05) by Fisher LSD *post-hoc* test. Correlation between FOS-induction in the SCN and in the retina were tested using a linear regression model: density ∼ (genotype*light) taking interaction effects into account. Results are expressed as mean ± SEM.

## Results

### Transcriptomic profiling of light-responsive genes in isolated retinal layers and the SCN reveals diverse responses

Given that both the retina and the SCN are sensitive to light, either directly or indirectly through retinal inputs, we hypothesized that genes differentially expressed in response to light may contribute to the phase-shifting responses of these clocks. To evaluate whether rods differentially shape transcriptional responses in the retina and the SCN, we profiled gene ecpression changes following an acute 530 nm light pulse designed to primarily target rods and known to shift both retinal clock and activity rhythms [34,37]. Bulk RNA sequencing was performed on whole SCN and on retinal layers isolated by laser microdissection from wild-type (WT) and *Opn4^-/-^::Trβ ^-/-^* (“rod-only”) mice exposed to a 530 nm monochromatic light pulse (10^14^ photons/cm²/s, 30 min) delivered at CT16 (Fig 1A). WT mice were euthanized either 30 min (n=3) or 2 h (n=3) after the onset of light stimulation. Consistent with previous findings in the SCN [30], light-evoked transcriptional changes were transient, with gene expressions at 2 h comparable to those of dark controls (DC) in WT mice (n=3). We therefore, focused on the 30-min time point to compare light-responsive transcriptomes between WT and “rod-only” mice. In WT mice, most differentially expressed genes (versus DC) were upregulated after 30 min of light exposure with 256 genes in the outer nuclear layer (ONL) of the retina, 34 in the inner nuclear layer (INL), 31 in the ganglion cell layer (GCL) and 17 in the SCN (Figs 1B-C, grey circles). Downregulated genes were comparatively few (30, 7, 4 and 3 genes respectively). In “rod-only” mice, the number of light-responsive genes was markedly reduced across all retinal layers and in the SCN, with very few genes showing increased expression compared to WT (Figs 1B-C, green circles). Interestingly, light-responsive genes in both genotypes displayed tissue and retinal layer-specific expression patterns (Fig 1C), with the exception of a shared cluster of immediate early genes, including *Fos*, *Junb*, *Nr4a1 and Btg2* that were induced in the INL, GCL and SCN (Figs 1C-D). As expected [62], *Fos* was downregulated in the ONL following light exposure, despite being robustly upregulated in inner retinal layers and the SCN.

**Figure 1.**
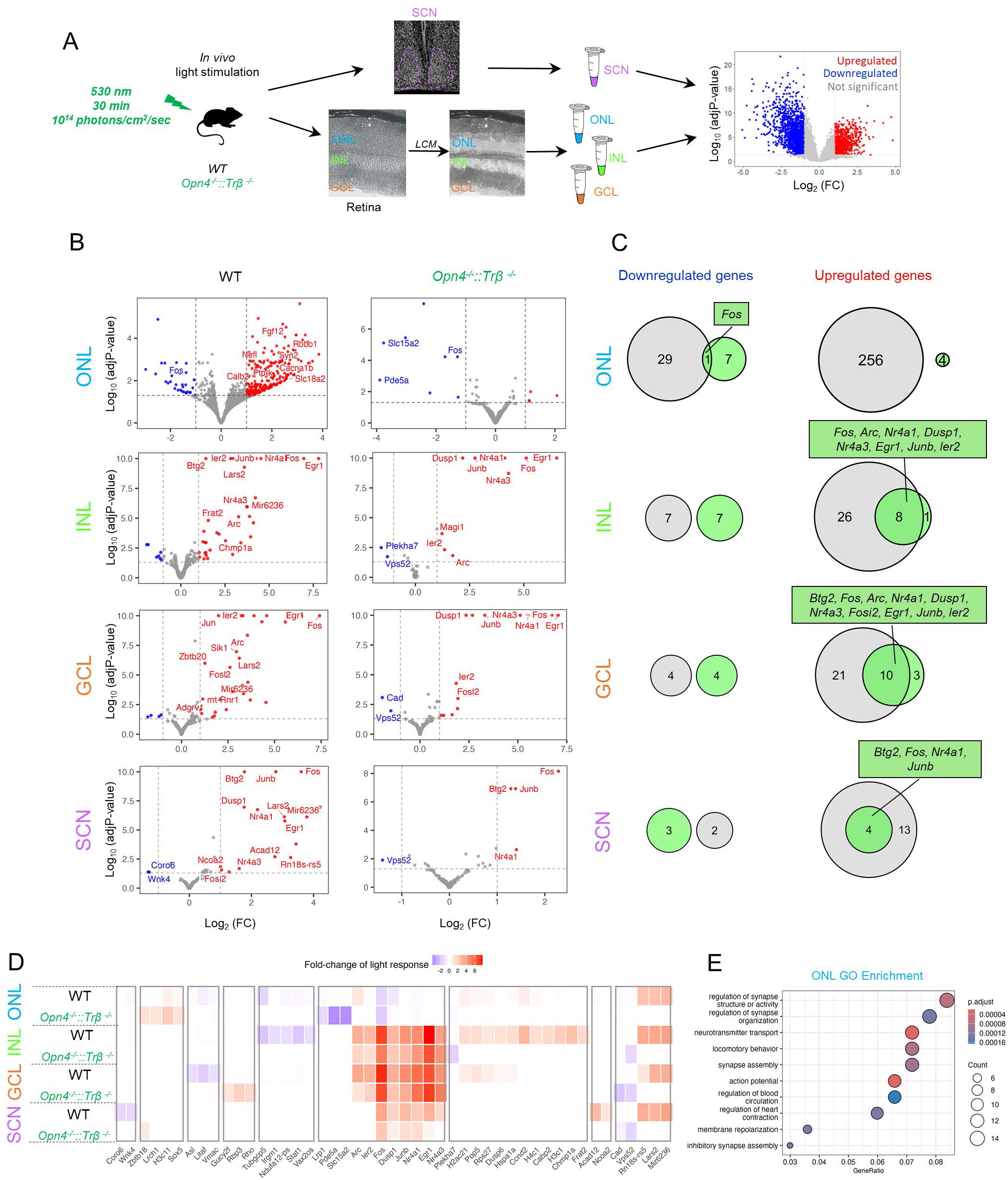
Transcriptomic profiles of light-responsive genes in isolated retinal layers and in the SCN following a monochromatic light pulse. A. Experimental workflow of 30-min of 530 nm light stimulation applied in wild-type (WT) and “rod-only” (*Opn4^-/-^::Trβ ^-/-^*) mice, followed by sampling of the SCN and eyes. Retinal layers (ONL, INL and GCL) were isolated using laser microdissection (LCM) and bulk RNA sequencing was performed (Dark controls or DC: WT, n=3 and *Opn4^-/-^::Trβ ^-/ -^*, n=3; for light-stimulated: WT: n=3 and *Opn4^-/-^::Trβ ^-/ -^,* n=3). B. Volcano plots showing differentially expressed genes in light-stimulated groups versus their respective DC in WT and “rod-only” mice. Significantly changed genes are defined by log_2_(FC) > 1 and p < 0.05. Red and blue dots respectively indicate upregulated or downregulated genes by light in ONL, INL or GCL or SCN. C. Venn diagrams showing specific and shared light-responsive genes between WT (grey circles) and “rod-only” (green circles) mice across ONL, INL, GCL and SCN. D. Heatmap clustering of light-responsive genes based on biological significance and similarity of light response between WT and “rod-only” mice in the ONL, INL, GCL and SCN. E. Gene Ontology (GO) enrichment analysis of light-responsive genes altered in the ONL of “rod-only” versus WT mice. GO enrichment was not performed for INL, GCL and SCN due to limited gene numbers.

To identify biological pathways altered in the “rod-only” mice compared with WT in response to light, we performed gene enrichment analysis using gene ontology (GO) terms and KEGG pathway analysis, focusing on the ONL which contained a sufficient and biological relevant number of light-responsive genes. In the ONL of “rod-only” mice compared to the ONL of light-stimulated WT, we observed downregulation of transcripts involved in synaptic signaling regulation, synapse organization, locomotor behavior, neurotransmitter transport, glutamate receptor and calcium signaling pathways such as *Calb2, Rab15, Scn1, Grm1, Grin2b* (Fig 1E). Interestingly, light-induced clock resetting in the SCN has been shown to depend on the transcription of immediate early genes [27,30–32]. Notably, *Egr1* and *Nr4a3* were no longer induced in the SCN of “rod-only” mice, suggesting a reduced photic signal reaching the master clock. Moreover, the light-induced expression of *Ncoa2*, a transcriptional coactivator of *Rorα* that positively regulates the circadian clock, was also attenuated in the SCN of “rod-only” mice.

### The absence of rods differentially impacts the light-induced FOS response in the retina and the SCN

To validate our transcriptomic data, we next examined FOS protein induction, a well-established marker of photic activation across tissues. FOS responses were assessed in WT, “rod-only” and additional genetic mouse models (Fig 2A) including *Nrl^-/-^* (“rodless”), *Rhoi75^Cre^::Cx36^flox/flox^* (lacking the gap junction protein connexin 36 (*Cx36*) specifically in rods, referred to *Rod^Cre^::Cx36^fl/fl^*) and *HRGP^Cre^::Cx36^flox/flox^* mice (lacking *Cx36* in cones, referred to *Cone^Cre^::Cx36^fl/f^* [52]. All mice were exposed to the same light stimulation (530 nm, 10^14^ photons/cm²/s, 30 min, CT16), previously shown to induce significant phase shifts of the retinal and SCN clocks [34]. FOS positive cells were quantified in response to light (Figs 2B-C) was quantified in INL and the GCL of the retina, as well as in the SCN of the same light-stimulated animals, and compared with their respective controls maintained in constant darkness (DC).

**Figure 2.**
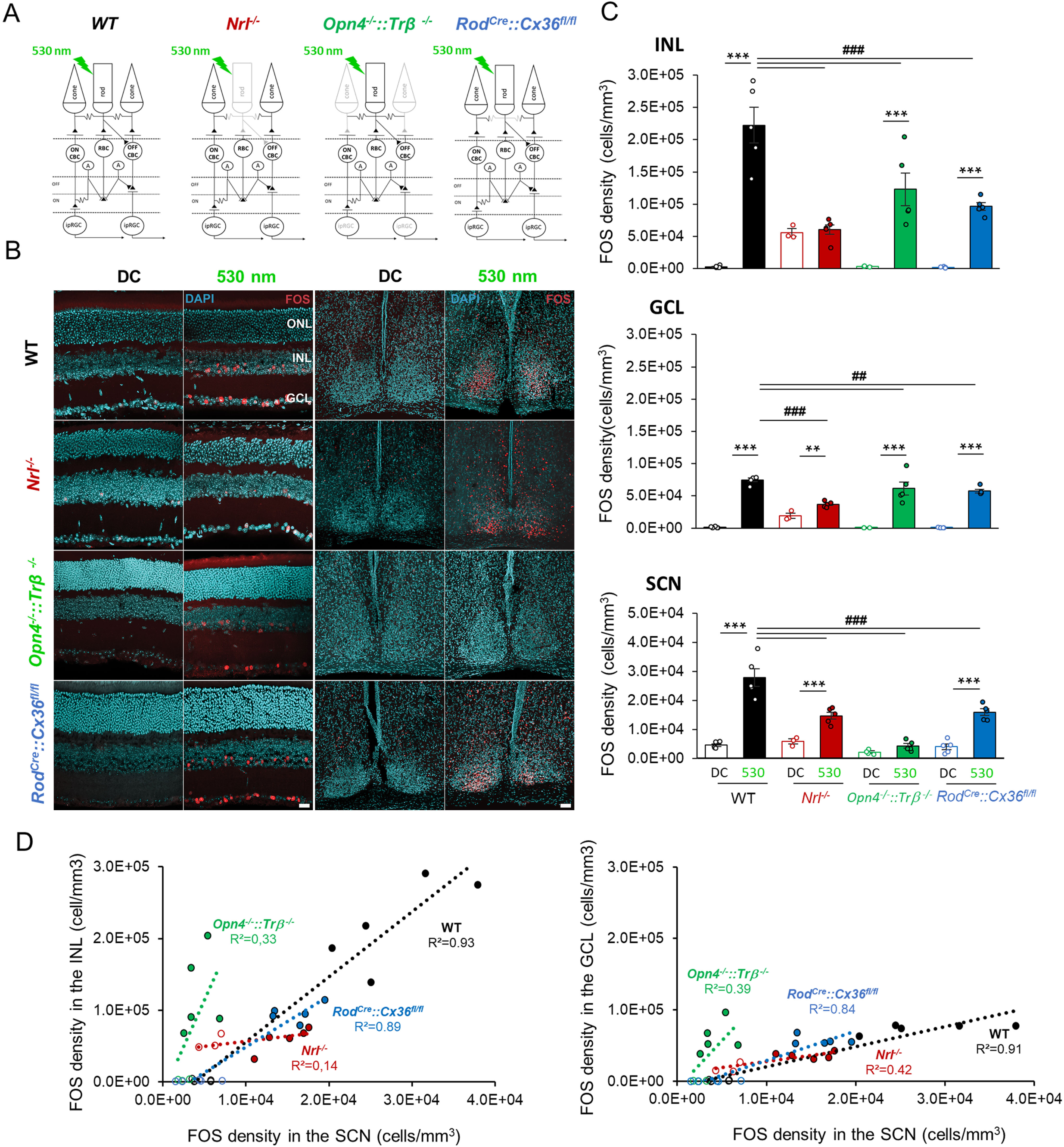
FOS induction in the retina and the SCN in response to a monochromatic pulse of light. **A.** Schematic representation of retinal circuitry originating from rods and cones in wild-type (WT), photoreceptor-deficient mice (*Nrl^-/-^* and *Opn4^-/-^::Trβ ^-/-^*) and in a mouse model with selective invalidation of the secondary rod pathway (*Rod^Cre^::Cx36^fl/fl^*). Only ipRGCs are indicated in the diagram at the level of the ganglion cell layer to simplify the network. RBC, rod bipolar cell; ON CBC, ON cone bipolar cell; OFF CBC, OFF cone bipolar cell; A, AII amacrine cells. **B.** Representative photomicrographs showing FOS-positive cells (red) in the retina and SCN of WT, *Nrl^-/-^, Opn4^-/-^::Trβ ^-/-^* and *Rod^Cre^::Cx36^fl/fl^* mice exposed to 530 nm light stimulation (30 min, 10^14^ photons/cm²/s) delivered at CT16, compared to dark controls (DC). ONL: outer nuclear layer; INL: inner nuclear layer; GCL: ganglion cell layer. Scale bar : 50 µm. **C.** Quantification of mean FOS cell densities in the INL, GCL and in the SCN in the same animals, across genotypes under dark (DC) and light-stimulated conditions. Bars represent mean ± SEM (for DC : WT: n=5; *Nrl^-/-^:* n=3; *Opn4^-/-^::Trβ ^-/ -^*: n=3; *Rod^Cre^::Cx36^fl/fl:^* n=5 ; for light-stimulated : WT: n=5; *Nrl^-/-^*: n=5; *Opn4^- /-^::Trβ ^-/ -^*: n=5; *Rod^Cre^::Cx36^fl/fl:^* n=5). Statistical differences with DC for each genotype are indicated by * with **: p ≤ 0.01 and ***: p ≤ 0.001 whereas statistical comparisons with light-stimulated WT animals are indicated by # with ## : p ≤ 0.01 and ###: p ≤ 0.001. **D.** Scatter plots showing correlations between FOS expression in the SCN and both the INL and GCL in response to light across different genotypes (empty and full dots represent individual data from DC and light-stimulated groups, respectively).

As expected [33,63], light significantly increases the number of FOS-positive cells in both the INL (222,284 ± 27,962 cells/mm^3^) and the GCL (74,532 ± 2,789 cells/mm^3^) of WT mice compared to their respective DC (INL: 2,502 ± 689 cells/mm^3^; GCL: 1,240 ± 423 cells/mm^3^, p < 0.001 for each layer; Fig 1C). Interestingly, in the “rodless” mice, light fails to induce FOS expression in the INL (60,324 ± 7,452 cells/mm^3^ compared to their respective DC: 55,906 ± 5,844 cells/mm^3^, p>0.05), but still elicits a significant increase in the GCL (light-stimulated: 36,820 cells/mm^3^; DC: 18,809 ± 4,197 cells/mm^3^, p<0.01) albeit to a lower extent than in WT light-stimulated retinas (p<0.001). This residual response likely reflects the contribution of ipRGCs and/or MW cones. In “rod-only” mice, the light-response capacity of rods alone is sufficient to trigger a substantial FOS response in both the INL (122,760 ± 25,617 cells/mm^3^; DC: 3,251 ± 552 cells/mm^3^, p<0.001) and the GCL (61,396 ± 9,934 cells/mm^3^; DC: 537 ± 224 cells/mm^3^, p<0.001), although in a diminished manner compared to WT retinas (p<0.001). Furthermore, the targeted deletion of *Cx36* in rods does not abolish FOS induction by light in either the INL (96,253 ± 5,820 cells/mm^3^; DC: 2,030 ± 393 cells/mm^3^, p<0.001) or the GCL (57,436 ± 2,740 cells/mm^3^; DC: 561 ± 159 cells/mm^3^, p<0.001). However as in the “rod-only” mice, the response remains attenuated in comparison to WT light-stimulated retinas (p<0.001). These data suggest that rods are the primary driver of the FOS response in both the INL and GCL, while melanopsin and MW cones contribute to GCL activation under photopic stimulation.

In the SCN of WT mice, 530 nm light stimulation robustly induces FOS expression (27,874 ± 3,081 cells/mm^3^ compared to DC: 4,543 ± 511 cells/mm^3^, p<0.001). Interestingly, unlike in the INL, FOS expression increases in response to light in the *“rodless”* mice (14,730 ± 1,233 cells/mm^3^ vs DC: 5,873 ± 816 cells/mm^3^, p<0.001) although with reduced levels compared to WT, indicating that non-rod photoreceptors can sustain SCN activation. However, in “rod-only” mice, light fails to significantly increase FOS expression in the SCN (4,307 ± 783 cells/mm^3^ vs 2,207 ± 381 cells/mm^3^ in DC; p>0.05). Moreover, both “rodless” and *Rod^Cre^::Cx36^fl/fl^* mice show a reduced FOS induction compared to WT (p<0.001), suggesting that rods contribute to SCN responsiveness, possibly via the secondary rod pathway under photopic conditions. This finding was further validated in the *Cone^Cre^::Cx36^fl/fl^* line (data not shown), in which the secondary rod pathway is lacking while indirect input from the primary rod pathway via the AII amacrine cells is still functional [52].

To examine the relationship between retinal and SCN responses to light, we applied a linear regression model to data from each animal, assessing the correlation between FOS induction in retinal layers and in the SCN (DC and light-stimulated; Fig. 2D). In WT and *Rod^Cre^::Cx36^fl/fl^* mice, FOS expression in the INL and GCL positively correlated with FOS induction in the SCN, indicating coordinated activation of both structures following 530 nm light stimulation (R^2^ ranging from 0.8-0.9). In contrast, *“*rodless*”* mice displayed an increased number of FOS positive cells in the SCN without a corresponding change in the retina, whereas *“*rod-only*”* mice exhibited robust retinal activation but no significant SCN response. These results reveal a clear dissociation between retina and SCN responses to high-irradiance 530 nm light, underscoring distinct mechanisms of photic processing in these two circadian clocks.

### Rods are required for the light-induced phase shift of the retinal clock but are not necessary for the SCN’s response at 530 nm photopic levels

Because the induction of FOS by light is not a direct indicator of circadian clock function, we next assessed retinal and SCN clock light sensitivity and the contribution of rod-mediated circuits, by measuring light-induced phase shifts *in vitro.* Retinal explants from *Nrl^-/-^*, *Opn4^-/-^::Trβ ^-/-^*, pan *Cx36^-/-^* and *Rod^Cre^::Cx36^fl/fl^* transgenic mice, all backcrossed onto the *Per2^Luc^* background were exposed to the same light stimulus used previously (520 nm, 10^14^ photons/cm²/s, 30 min, CT16; Figs 3A-B). In parallel, we quantified light-induced phase shifts of locomotor activity rhythms in these genotypes (Figs 3C-D).

**Figure 3.**
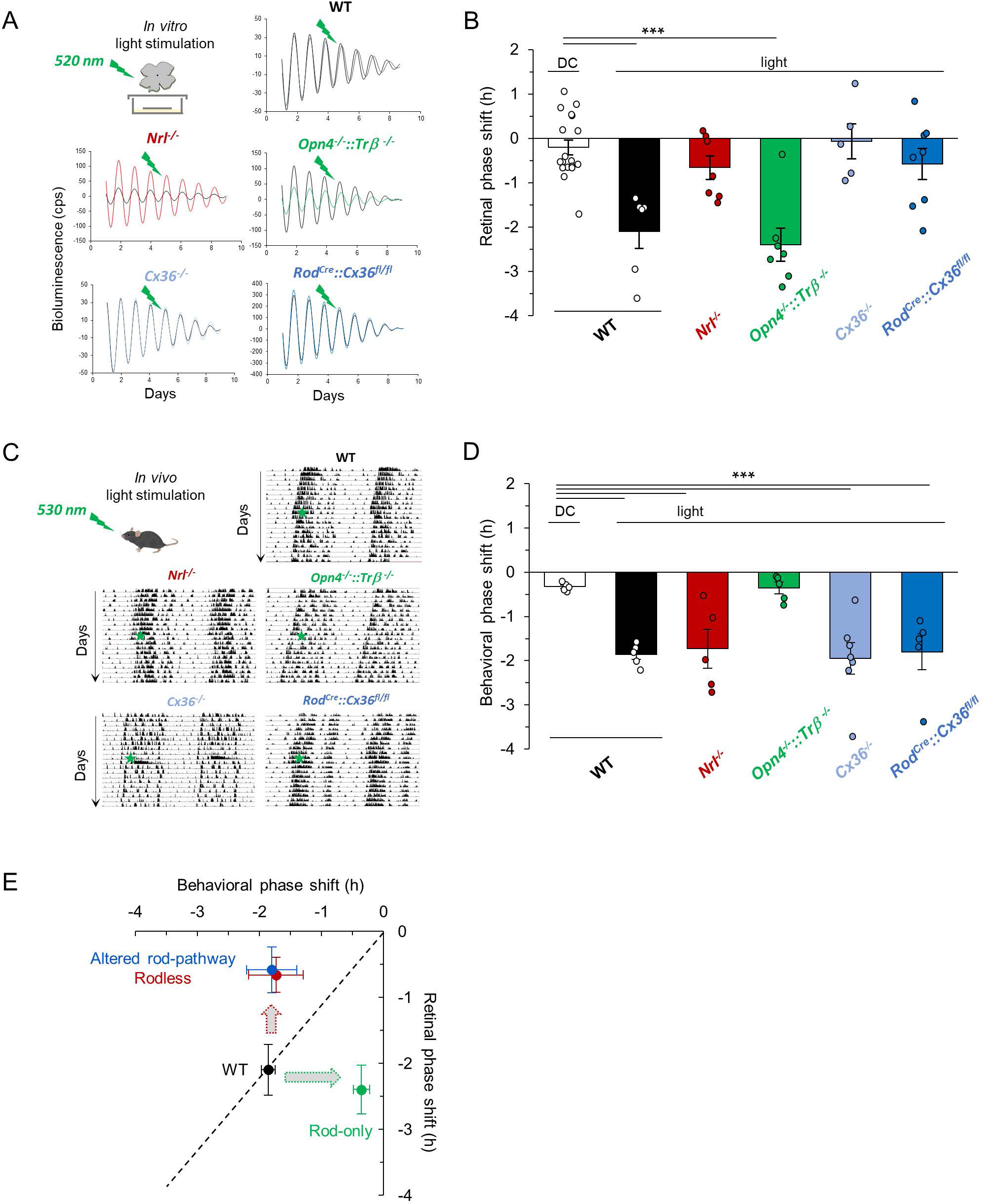
Light-induced phase-shifting of the retinal clock requires rods, in contrast to the SCN. **A.** Representative PER2::Luc bioluminescence traces from retinal explants of wild-type *Per2^Luc^* (WT), photoreceptor-deficient (*Nrl^-/-^* and *Opn4^-/-^::Trβ ^-/-^*) and *Rod^Cre^::Cx36^fl/fl^* mice exposed to a light stimulation (colored lines) compared to DC (black line). **B.** Mean light-induced phase shift of the retinal clock in response to a light stimulation (520 nm, 30 min, 10^14^ photons/cm²/s) applied at CT16. Bars represent mean ± SEM (DC: n=18; WT: n=6: *Nrl^-/-^:* n=7; *Opn4^-/-^::Trβ ^-/-^* : n=7; *Cx36^-/-^ : n=5 ; Rod^Cre^::Cx36^fl/fl:^* n= 8). **C.** Representative actograms of locomotor activity in WT, *Nrl^-/-^, Opn4^-/-^::Trβ ^-/-^* and *Rod^Cre^::Cx36^fl/fl^* mice exposed to a light pulse (530 nm, 30 min, 10^14^ photons/cm²/s) at CT16 (green star). **D.** Mean light-induced phase shift of the rhythm of locomotor activity. Bars represent mean ± SEM (DC: n=5 WT: n=5: *Nrl^-/-^:* n=5; *Opn4^-/-^::Trβ ^-/ -^*: n=5; *Cx36^-/-^ : n=7 ; Rod^Cre^::Cx36^fl/fl:^* n= 5). Statistical differences with the DC are indicated by ***: p ≤ 0.001. E. Relationship between mean light-induced phase shifts (± SEM) in the retina and SCN for WT (black dot), “rodless” (red dot), “rod-only” (green dot) and *Rod^Cre^::Cx36^fl/fl^* (blue dot) mice.

Consistent with earlier findings [34], we confirmed that rods are required for the light-induced phase shift of the retinal clock at high irradiance (Fig 3B). In *“*rodless” retinal explants, no significant phase shift in response to light is observed (−0.66 ± 0.26h; DC: - 0.19 ± 0.14h, p>0.05), whereas a significant phase delay is detected in retinal explants from “rod-only” mice (−2.40 ± 0.37h, p<0.001 compared to DC), comparable to that observed in WT explants (−2.10 ± 0.38h).

To identify the circuits responsible for rod-mediated light signals that induce phase shift of the retinal clock, we first examined the role of gap junction communication via *Cx36* (Figs 3A-B). We used pan *Cx36^-/-^* mice which lack gap junctions between AII amacrine cells and ON-CBCs in the INL, thereby disrupting the primary rod pathway input. Additionally, these mice lack the secondary rod pathway due to the absence of rod/cone coupling. In this genotype, light failed to induce a phase shift (−0.06 ± 0.39h, p>0.05 compared to DC; Fig 3B) in these mice. To specifically assess the contribution of the secondary rod pathway, we analyzed retinal explants from *Rod^Cre^::Cx36^fl/fl^* mice, which also showed no significant light-induced phase shift (−0.58 ± 0.35h, p>0.05 compared to DC). Together, these results indicate that the secondary rod pathway, operating through cones, is sufficient to mediate the light response of the retinal clock under photopic conditions.

To further investigate the dissociation between rod-mediated light-responses in the retina and the SCN, mirroring the divergenve observed in FOS responses, we measured light-induced phase shifts of locomotor activity rhythms in the same genotypes. Phase shifts were similar across WT (−1.86 ±0.11h), “rodless” (−1.75 ± 0.42h) and *Rod^Cre^::Cx36^fl/fl^* (−1.8 ± 0.40h) compared to DC (−0.33 ± 0.04h ; p<0.001 for all comparisons; Fig 3D), indicating that neither rods nor their electrical coupling to cones or other rods is essential for shifting the phase of the central clock. Conversely, no phase shift was observed in “rod-only” mice (−0.36 ± 0.13h; p>0.05), indicating that rods alone are insufficient to elicit a behavioral phase shift under our light conditions. To compare light response dynamics between the retinal and SCN clocks across genotypes, we plotted the average behavioral phase shift against the average retinal phase shift for each genotype (Fig 3E). In WT mice, 520-530 nm light stimulation caused phase shifts in both clocks. However, loss of rods or disruption of rod/cone electrical coupling abolished the retinal clock’s phase shift while leaving the behavioral phase shift unchanged. Conversely, when only rods were present, light induced a robust retinal clock responses, but failed to elicit a behavioral phase shift.

### Rods use the OFF-cone bipolar cell pathway to evoke a light-induced phase shift of the retinal clock

Our data in Fig 3 strongly suggest that rods rely on the secondary rod pathway to mediate light-induced phase shift of the retinal clock. However, the genetic strategy used does not differentiate between the ON and OFF component of the secondary rod pathway, nor do it selectively isolate the tertiary rod pathway. To overcome these limitations, we combined genetic mouse models with a pharmacological strategy that selectively blocks individual rod pathways (Fig 4A). The ON response to light results from the interruption of glutamate transmission from photoreceptors onto ON bipolar cells. To investigate the role of ON pathways to retinal clock phase shifting, we first supplemented the culture medium containing wild-type *Per2^Luc^* retinal explants with L-AP4 (l-(+)-2-amino-4- phosphonobutyric acid), a selective agonist of metabotropic glutamate receptor 6 (Fig 4A). L-AP4 blocks glutamatergic signaling from rods to RBCs and from cones to ON-CBCs [46]. In the presence of L-AP4, 520 nm light stimulation induced a robust phase delay of the retinal clock (−2.88 ±0.49h, p<0.001) comparable to the delay observed without L-AP4 supplementation (−2.10 ± 0.38h, p>0.05). These results indicate that the ON pathways, whether via RBCs and AII amacrine cells (primary pathway) or *via* cones to ON-CBCs (ON secondary pathway) are not required for light-induced phase shifting of the retinal clock. We next treated retinal explants from *Rod^Cre^::Cx36^fl/fl^::Per2^Luc^* mice with L-AP4. In this case, only the tertiary rod pathway remain functional. Under these conditions, no significant phase shift was observed (0.01 ± 0.31h, p>0.05 compared to DC) suggesting that the tertiary rod pathway alone is insufficient to drive retinal clock resetting. Finally, we treated WT *Per2^Luc^* retinal explants with a combination of L-AP4 and UBP310 ((S)-1-(2-Amino-2-carboxyethyl)- 3- (2-carboxy-thiophene-3-yl-methyl)- 5 -methylpyrimidine-2,4-dione), an antagonist of kainate receptors on OFF-CBCs that block the transmission from both rods and cones to OFF-CBCs. No phase shift was observed (0.26 ±0.34h, p>0.05 compared to DC). Collectively, these results demonstrate that the secondary rod pathway operating through OFF-CBCs is essential for mediating the retinal clock’s response to light.

**Figure 4.**
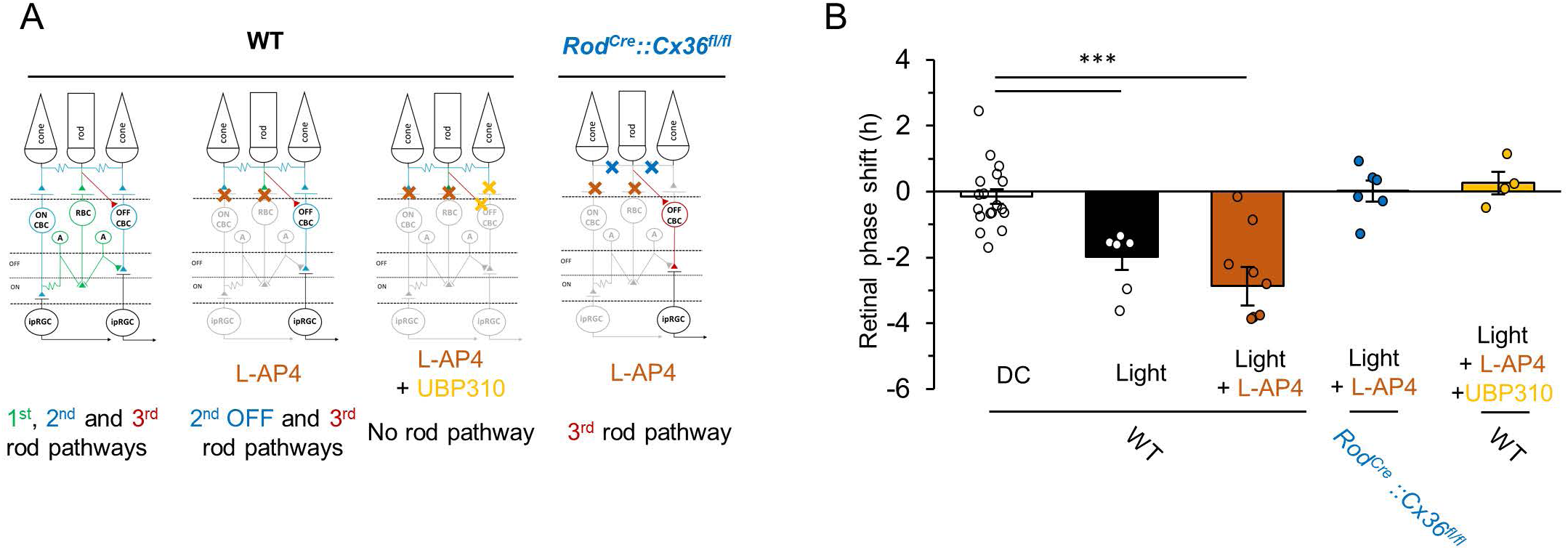
Rods shifts the phase of the retinal clock through the secondary rod pathway involving OFF cone bipolar cells. **A.** Schematic representation of the primary, secondary and tertiary rod pathways and their pharmacological blockade by L-AP4 (brown cross) or a combination of L-AP4 and UBP310 (yellow cross). mGluR6 receptors expressed on ON cone bipolar cells (ON-CBCs) and rod bipolar cells (RBCs) are sensitive to L-AP4. In the WT *Per2^Luc^* mice, L-AP4 blocks the primary and secondary ON pathways leaving only the secondary OFF and tertiary pathways functionals. In *Rod^Cre^::Cx36^fl/fl^ ::Per2^Luc^* explants treated with L-AP4, only the tertiary pathway remains functional. The combined supplementation of L-AP4 and UBP310 blocks all rod-driven signals. **B.** Mean phase shifts of the retinal clock following 530 nm light stimulation, measured in the absence or presence of L-AP4 (n=8) or L-AP4 plus UBP310 in WT (n=4) and *Rod^Cre^::Cx36^fl/fl^ ::Per2^Luc^* (n= 6) explants. Bars represent mean ± SEM. Statistical differences with DC are indicated by ***: p ≤ 0.001.

## Discussion

This study investigates the molecular basis of light-evoked responses in the retina and the SCN by combining transcriptomics with FOS immunodetection to reveal how each tissue engage distinct photoreceptor-driven signaling pathways. Although heterogeneity in SCN neuronal light responsiveness is well documented [64,65], our work reveals, for the first time, layer-specific transcriptional activation in the retina following photic stimulation. In WT mice, 530 nm light stimulation at CT16, previously shown to shift the phase of the retinal clock [34], triggered robust gene expression, particularly in the ONL, consistent with intact photoreceptor function. In contrast, in “rod-only” mice lacking MW cones and melanopsin, transcriptional responses are markedly reduced across retinal layers and in the SCN, demonstrating that rods alone cannot sustain photic gene activation under these photopic conditions. Gene ontology analysis of downregulated ONL transcripts in “rod-only” compared to WT mice, revealed impaired signaling pathways, especially those associated with synaptic transmission, further underscoring the requirement of MW cones and melanopsin for initiating a full light-responsive transcriptional program. The spectral property of the stimulus likely contribute to these deficits, as 530 nm light lies outside the optimal sensitivity range of SW cones and neuropsin (OPN5), both UV-sensitive opsins [66]. Although OPN5 has been proposed to mediate retinal clock entrainment (Buhr et al., 2015), its activation under these conditions is expected to be minimal. Nevertheless, residual induction of immediate early genes (*Fos, Junb, Nr4a1, Nr4a3, Btg2, Egr1, Ier2*) in the INL and GCL of “rod-only” retinas suggests that rod signaling may be sufficient to shift the retinal clock (Figs. 3B–C). In the SCN, *Fos*, *Junb*, and *Nr4a1* remained light-inducible in “rod-only” mice, contrasting with the profound reduction in FOS induction observed in *Opn4^-/-^* animals (∼92% decrease; [67]. Despite this preserved immediate early gene activation, no behavioral phase shift occurred, indicating that rod input is too weak to initiate the full transcriptional cascade required for SCN clock resetting in the absence of melanopsin [5,9,68]. Notably, *Egr1* failed to respond to light in the “rod-only” SCN. However, given that *Egr1^-/-^* mice display normal behavioral phase shifts in response to bright light [69,70], *Egr1* is unlikely to be essential for this process. Instead, its loss may reflect broader alterations in glutamatergic signaling, as EGR1 regulates NMDA-dependent transcription of PSD-95 [71,72] and influences AMPA receptor trafficking, mechanisms known to contribute to photic phase shifting [73–75].

We then assessed FOS protein, a marker that is consistently light-induced in the retina and SCN of WT and “rod-only” mice, and extended this analysis to additional relevant mouse models. Strikingly, in “rodless” mice, which lack rods but retain functional cones capable of signaling through RBCs [50,76,77], light failed to induce FOS in the INL and produced a reduced response in the GCL, consistent with previous observations [78]. In contrast, “rod-only” mice displayed persistent though attenuated FOS induction in both layers, highlighting the capacity of rods to drive retinal activation, particularly within the INL. At the level of the SCN, FOS induction was diminished in “rodless” mice and absent in “rod-only” mice, reinforcing the critical roles of MW cones and melanopsin in mediating central photic signaling [3,6,79]. Because, rod-cone gap junctions constitute the predominant coupling mechanism among photoreceptors [52], we next examined *Rod^Cre^::Cx36^fl/fl^* mice to determine whether rod-derived signals are transmitted through this pathway to influence retinal clock resetting. These mice exhibited a reduced light-induced FOS response similar to that of “rod-only” mice. A similar phenotype was observed in *Cone^Cre^::Cx36^fl/fl^* mice, which closely parallel the *Rod^Cre^::Cx36^fl/fl^* model [80], suggesting that residual FOS induction arises from the transmission of rod signals to cones via the secondary rod pathway.

Although light-induced FOS expression provides a reliable readout of neural activation, it does not directly reflect clock resetting by light. This is exemplified in *Rod^Cre^::Cx36^fl/fl^* mice where light induces FOS in the retina despite having an effect on the retinal clock. At 530 nm, light fails to shift the phase of the retinal clock, yet it elicits robust behavioral phase shifts in “rodless” mice. Conversely, “rod-only” mice display retinal phase delays, without corresponding behavioral effects indicating that rods can reset retinal but are unsufficient to drive SCN entrainment. This lack of behavioral phase shift has also been observed at lower irradiance levels “rod-only” mice [34] further supporting the essential contribution of melanopsin and MW cones. Interestingly, melanopsin alone can drive light-induced phase shifts of the retinal clock during the first post-natal week before rod functionality emerges [81], underscoring the differential contributions of rods and melanopsin at different developmental stages. A comparable “rod-only” model generated by expressing an attenuated diphtheria toxin A subunit under a cone-specific promoter in a melanopsin null background [2], similarly failed to exhibit behavioral phase shifts despite intact photic entrainment under photopic and scotopic conditions. Consistent with this, retinal phase shifts are abolished in both pan *Cx36^-/-^::Per2^Luc^* and *Rod^Cre^::Cx36^fl/fl^::Per2^Luc^* mice, in which the secondary ON and OFF rod pathways are selectively disrupted. Despite this loss, behavioral phase shifts by light remain intact, demonstrating a functional dissociation between retinal and SCN entrainment mechanisms and confirming that the secondary rod pathways via *Cx36* are dispensable for SCN-mediated behavioral responses. These findings also indicate that retinal clock resetting does not require the primary rod pathway, which depends on *Cx36*-mediated gap junctions between AII amacrine cells and between AII cells and ON CBCs. However, the absence of *Cx36* in the inner retina significantly lengthens the endogenous period of the retinal clock, which consists of a network of multiple coupled cellular clocks [82,83]. Thus, while *Cx36* is not essential for sustaining PER2::Luc rhythms, it plays a critical role in coordinating the temporal coherence of this multicellular oscillator network.

The unexpected contribution of rods at bright light intensities, within the sensitivity range of cones, has been proposed previously for non-visual responses to light, such as the pupillary light reflex (PLR) and retinal responses involving dopaminergic amacrine cells [2,6,84,85]. In contrast to our findings, Beier et al. (2022) recently reported that under photopic light levels, rods can drive both pupil constriction and behavioral circadian photoentrainment via both the primary and secondary ON pathways [86]. These divergent conclusions likely reflect substancial differences in the light parameters and physiological outputs assessed: we employed long-duration, high-irradiance monochromatic light stimulation (30 min, 10^14^ photons/cm²/s, 520-530 nm) to evaluate phase-shifting of both retinal and SCN clocks. By contrast, Beier *et al.* used brief, lower-intensity white light pulses (30 s, white light at 100 lux) to assess the PLR. Furthermore, PLR measurements were conducted between ZT4 and ZT12, whereas our phase shifting experiments were performed at CT 16.

To determine which rod pathways mediate light-induced retinal phase shifting, we pharmacologically inhibited the primary rod input to RBCs and the cone input to ON-CBCs in *Per2^Luc^* retinal explants. This medium supplementation did not alter the phase shifting response in WT explants but abolished it in *Rod^Cre^::Cx36^fl/fl^::Per2^Luc^* cultures. These findings suggest that light adjusts the retinal clock phase primarily through the secondary OFF pathway, whereas the tertiary rod pathway does not contribute to this response. The OFF-cone pathway, specialized for detecting decreases in light intensity becomes particularly active during the transition from light to darkness [87]. Under natural conditions, the retina experiences sustained illumination throughout the day. Thus, recruitment of the OFF-CBC circuit, together with the low light sensitivity of the retinal clock ([34], present study) may enhance the clock’s ability to detect gradual dimming and encode dark signals following a light pulse, a potent synchronizing cue in mice [88]. Light regulation of the retinal clock may therefore be fine-tuned through the comparatively more developed OFF pathway [89]. The high density of ribbon synapses on OFF-CBCs axons contacting RGCs likely supports a robust local regulatory network capable of precise modulation of synaptic neurotransmitter release [44]. In parallel, dopamine signaling may provide an additional modulatory layer. Under similar photopic conditions than the current study, dopamine release has been reported to be rod-driven [84] and to reset the retinal clock through activation of D1 dopamine receptors [90]. Via these same receptors, dopamine decreases glycinergic inhibition onto OFF-CBCs, thereby adjusting their sensitivity to changes in light intensity [87,91,92].

## Acknowledgments

We acknowledged C Coutanson for her technical help and Y Dusabyinema and M Fackeuere at the Plateforme Sequencage IGFL (PSI) for the RNA-seq analysis.

## Author Contributions

A.J. and O.D-B. conceived the study. A.J. and H.C. performed the experiments; C.S. performed RNA extraction; A.D-S., K.T., B.A. and K.P. analysed RNA-seq data. A.J., H.C., N.H., MP. F-S and O.D-B. analysed data. C.R. generously provided the *Cx36^fl/fl^, Cx36^-/-^ Rod^Cre^* and *Cone^Cre^* mouse lines. A.J. and O.D-B. wrote the manuscript. All authors were involved in reviewing and editing.

## Data and Materials Availability

Further information and requests for resources and reagents should be directed to and will be fulfilled by the lead contact, Ouria Dkhissi-Benyahya.

